# Variable expression quantitative trait loci analysis of breast cancer risk variants

**DOI:** 10.1101/2020.03.23.004366

**Authors:** George Wiggins, Michael A Black, Anita Dunbier, Tony R Merriman, John Pearson, Logan C Walker

## Abstract

Genome wide association studies (GWAS) have identified more than 180 variants associated with breast cancer risk, however the underlying functional mechanisms and biological pathways which confer disease susceptibility remain largely unknown. As gene expression traits are under genetic regulation we hypothesise that differences in gene expression variability may identify causal breast cancer susceptibility genes. We performed variable expression quantitative trait loci (veQTL) analysis using tissue-specific expression data from the Genotype-Tissue Expression (GTEx) Common Fund Project. veQTL analysis identified 70 associations (*p* < 5×10^−8^) consisting of 60 genes and 27 breast cancer risk variants, including 55 veQTL that were observed in breast tissue only. Pathway analysis of genes associated with breast-specific veQTL revealed an enrichment of four genes (*CYP11B1, CYP17A1 HSD3B2* and *STAR*) involved in the C21-steroidal biosynthesis pathway that converts cholesterol to breast-related hormones (e.g. oestrogen). Each of these four genes were significantly more variable in individuals homozygous for rs11075995 (A/A) breast cancer risk allele located in the *FTO* gene, which encodes an RNA demethylase. The A/A allele was also found associated with reduced expression of *FTO*, suggesting an epi-transcriptomic mechanism may underlie the dysregulation of genes involved in hormonal biosynthesis leading to an increased risk of breast cancer. These findings provide evidence that genetic variants govern high levels of expression variance in breast tissue, thus building a more comprehensive insight into the underlying biology of breast cancer risk loci.

## INTRODUCTION

Genome wide association studies (GWAS) in breast cancer have identified more than 180 common risk variants^1–3^, however the causal genes and biological mechanisms which confer disease susceptibility remain largely unknown. Risk variants are often located in non-coding regions making it difficult to determine pathogenic pathways. Approximately 700 potential gene targets of breast cancer risk variants have been identified using analytical methods that employ genomic data from chromatin interactions, enhancer–promoter correlations, transcription binding, topologically associated domains and gene expression^1,3^.

Gene expression traits are under genetic regulation and the heritability of differences in genotypes have been extensively described^4^. For example, identification of expression quantitative trait loci (eQTL) has been a key approach for investigating tissue-specific effects of breast cancer risk variants under the hypothesis that non-breast tissue may be involved in breast cancer risk^5^. Gene expression patterns are often explored assuming genetic control of mean expression level, however the variability of gene expression is also genetically controlled^6–9^. Just as differences in expression means have been associated with genotype so too differences in expression variability can be associated with genotype.

Gene expression variability has been described in a wide range of organisms including prokaryotes^10^, yeast^6,7^ and complex multicellular organisms^11–13^. Furthermore, gene expression variability had been shown to be important in early human development^14^, schizophrenia^15^ and cancer subtypes^12,13^. The effects of genetic variation on gene expression variability has been recently described in human derived lymphoblastoid cell lines from HapMap individuals^8^ and in the TwinsUK cohort^16,17^.

Breast cancer risk variants associated with eQTL, based on mean gene expression, have been investigated in both breast tissue (tumour and normal), and non-breast tissue^5,18–20^. However, the mechanisms underlying breast cancer risk for the majority of variants remains to be uncovered. Here, we demonstrate variable expression quantitative trait loci (veQTL) as a method for testing the association of variants with gene expression variability. We performed veQTL analysis on 181 variants that have been previously associated with breast cancer risk and identified 60 new candidate genes and pathways associated with 27 breast cancer risk variants.

## METHODS

### Data acquisition and processing

Genotype and expression data were acquired through the database of Genotypes and Phenotypes (dbGaP) and the Genotype-Tissue Expression (GTEx) Common Fund Project (release version phs000424.v7.p2.) under the project title “Identification of variable expression quantitative trait loci that are associated with cancer risk”. Datasets from breast, ovarian, lung and kidney tissue used in this study were obtained through the dbGaP approval number 17463.

Genotype data from 635 individuals acquired through GTEx were converted to chromosome-specific matrices, where the genotypes were numbered by the minor allele count. For tissue specific analysis, only genotypes from individuals with tissue expression data in a given tissue (e.g. breast, kidney, ovary and lung) were retained. Genotypes were filtered so that only bi-allelic genotypes of at least 10 subjects with two or more genotypes (AA, Aa, aa) were retained.

Normalised Reads Per Kilobase of transcript, per Million mapped reads (RPKM) counts for 56,203 unique Ensembl (https://www.ensembl.org/) gene ids were split into tissue-specific datasets. For each dataset, only transcripts with RPKM > 0.1 in at least 10 samples were retained. Subjects with multiple tissue-specific samples were collapsed by calculating the average RPKM values. Linear regression models were used to correct expression data for age and sex as covariates.

### veQTL and eQTL analysis

Tissue specific veQTL were mapped for breast cancer risk variants that passed the filtering criteria (Supplementary Table 1). For each gene, veQTL were mapped by testing for equal variance among individuals of different genotypes using the Brown-Forsythe method^21^. A custom R script (https://github.com/jfpuoc/veQTL) was used to calculate Brown–Forsythe test-statistics (*W*, equation 1) on each genotype and all transcripts. For a response variable *y* in *j* groups, transformed to the median absolute deviation *Z*_*ij*_ = |*y*_*ij*_ – *y*_*j.*_| where *y*_*j.*_ is the median in group *j*, then *W* is defined by:

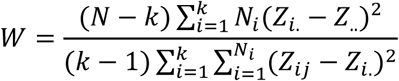

Equation 1. Brown-Forsythe test statistic

Where *N* is the number of samples, *k* is the number of different genotypes (2 or 3), *N*_*i*_ is the number of samples in group *i, Z*_*i*_ is the mean of the absolute deviation from the medians for group *i* and *Z*_.._ is the mean of the absolute deviations from all samples from their respective group medians. The resulting *W* statistics follows the F-distribution with degrees of freedom *df1* = *k – 1* and *df2* = *N – k* ^21^.

veQTL analysis was performed using the residuals of the linear model correcting for age and sex, and the genotypes that met the filtering criteria. In instances where two genotypes were observed in more than 10 samples, and the third genotype was observed in less than 10 samples, the test statistic was only computed between groups with at least 10 samples.

Tissue-specific eQTL analysis was performed in the same four tissue datasets used for veQTL. The ultra-rapid MatrixEQTL package in R was used to calculate *p-*values for variant-gene pairs using a linear regression model and correcting for age and sex as covariates^22^.

We limited proposed breast cancer susceptibility genes to those that had: i) significant (*p* < 5.0×10^−8^) gene expression variability associated with a breast cancer risk variants, ii) the significant veQTL association was only observed in breast tissue and iii) the gene was only associated with a change in expression variability (i.e. veQTL) and not change in mean expression (i.e. eQTL).

### Pathway enrichment analysis

Genes identified with altered expression by either veQTL or eQTL analysis were annotated using their entrez identifier. Pathway analysis was performed using the R packages clusterProfiler and DOSE^23,24^. Each candidate gene list was compared to the background transcriptome for over representation of genes in pathways annotated by GO terms.

## RESULTS

### Identification of veQTLs and eQTLs

The GTEx dataset comprises 635 genotyped samples, of which tissue samples from normal breast (n=255), lung (n=387), kidney (n=41) and ovary (n=123) were used. A significant proportion of breast cancer risk variants are predicted to alter expression of cancer susceptibility gene(s) in breast tissue. To identify veQTL that specifically increase risk in breast tissue, even if the genes in the veQTL are ubiquitously expressed in multiple tissues, we only considered veQTL in breast tissue that were not identified in other tissue types. These assumptions, would however miss breast cancer susceptibility genes whose expression variability is tolerated in other tissue but not breast.

RNA-sequencing and genotype data were split into tissue-specific datasets and filtered to remove low frequency genotypes and genes with low expression. After pre-processing 33059, 29522, 25026 and 35137 transcripts were retained for the breast, ovary, kidney and lung, respectively.

Large genome-wide association studies (GWAS) have identified variants associated with breast cancer risk or subtype specific breast risk. In total we identified 181 breast cancer risk variants in the literature (Supplementary Table 1), of which 152, 148, 106 and 152 breast cancer risk variants were retained after filtering non-biallelic and genotypes with few minor alleles (see methods) for the breast, ovary, kidney and lung datasets, respectively (Figure 1, Supplementary Table 1).

**Figure. 1.**
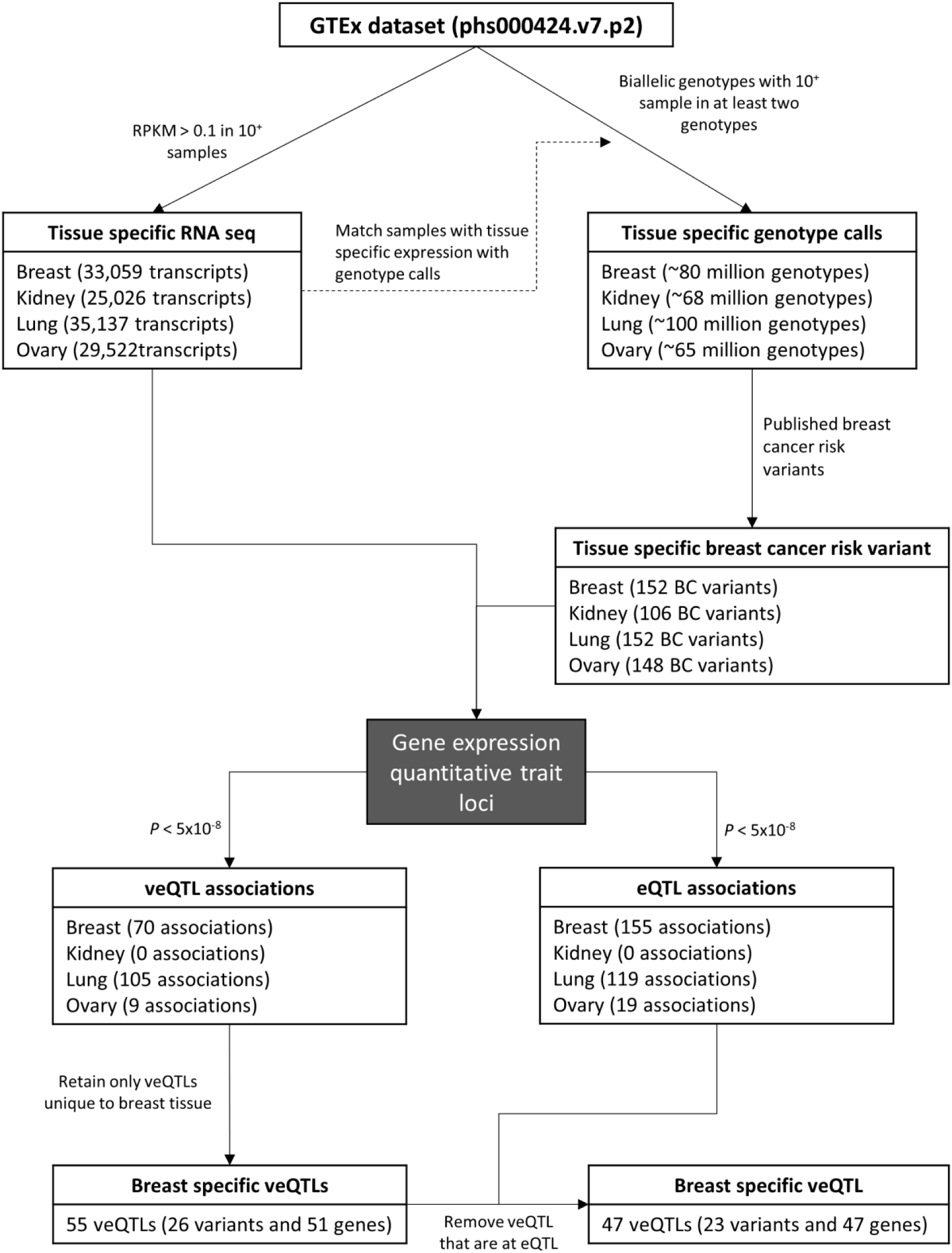
Schematic of study rationale to identify veQTL and eQTL in four different tissue types

We tested for associations between breast cancer risk variants and gene expression variability, correcting for sex and age, in four tissues. The risk variants were significantly (*p* < 5×10^−8^) associated with veQTL in the breast (70), ovary (9) and lung (109) (Table 1). No significant associations were observed in the kidney analysis. By comparison, the number of observed eQTL in breast (155), ovary (19) and lung (123) were greater, similarly there were no significant kidney eQTL. The majority of veQTL and eQTL associations were *trans* and acted over distances greater than 1 Mb or between chromosomes. Only 2/70, 5/109 and 2/9 significant association were *cis-*veQTL (+/- 1 Mb) in the breast, lung and ovary, respectively. A greater proportion of eQTL were observed in *cis* compared to veQTL, with approximately 5% of veQTL and 13% eQTL acting in *cis*.

**Table 1.** Significant veQTL and eQTL breast cancer variants and associated genes for each tissue

|  | veQTL |  |  | eQTL |  |  |
| --- | --- | --- | --- | --- | --- | --- |
|  | Variants | Genes | veQTL (*cis*) | Variants | Genes | eQTL (*cis*) |
| Breast | 27 | 60 | 70 (2) | 16 | 139 | 155 (18) |
| Kidney | 0 | 0 | 0 | 0 | 0 | 0 |
| Lung | 28 | 81 | 109 (5) | 24 | 101 | 123 (17) |
| Ovary | 5 | 9 | 9 (2) | 7 | 19 | 19 (4) |

### Classes of veQTLs

By assessing expression values associated with each genotype across the four different tissues, we observed three classes of veQTL (Figure 2). Class I resembled a homozygous recessive phenotype, where the presence of two minor alleles was associated with altered gene expression variability. Class II showed a dominant phenotype where the dosage of the minor allele correlated with the change in expression variability. Class III resembled a heterozygous phenotype where the presence of two different alleles altered gene expression variability. Significant breast veQTL were largely Class I homozygous recessive (56%), (Figure 2), while the majority (9/11) of Class II veQTL were also eQTL. In total, 21 veQTL (30%) were also eQTL. Seven breast cancer variants with significant veQTLs had no samples homozygous for the minor allele and were unable to be classified. However, all seven variants had more gene expression variability in heterozygous samples, thus ruling out a Class I veQTL.

**Figure 2.**
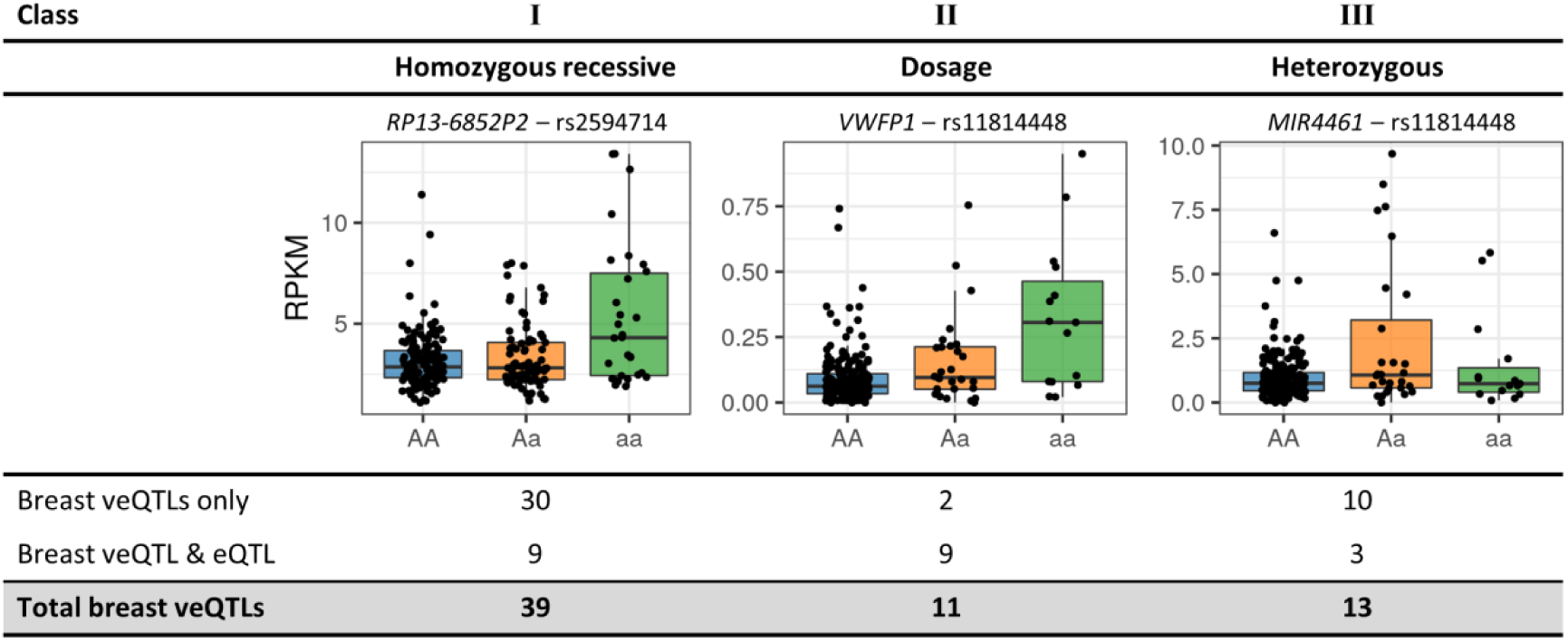
Characteristics of gene expression variability in veQTLs. Three class of veQTLs were observed with respect to the minor allele. Significant breast veQTLs were represented in all three classes with the majority (39/70) class I.

### Comparison of veQTL and eQTL

To estimate biases in dataset specific veQTL analysis quantile-quantile plots (q-q plots) were generated and genomic factors estimated for each tissue (Figure 3a). No substantial genomic inflation (λ < 1.1) was observed for the veQTL analysis in the breast, lung or ovary (λ ranged 1.00-1.05). However, a larger genomic inflation factor of 1.15 was observed for kidney tissue, implying a small underlying bias in the analysis (Figure 3a).

**Figure 3.**
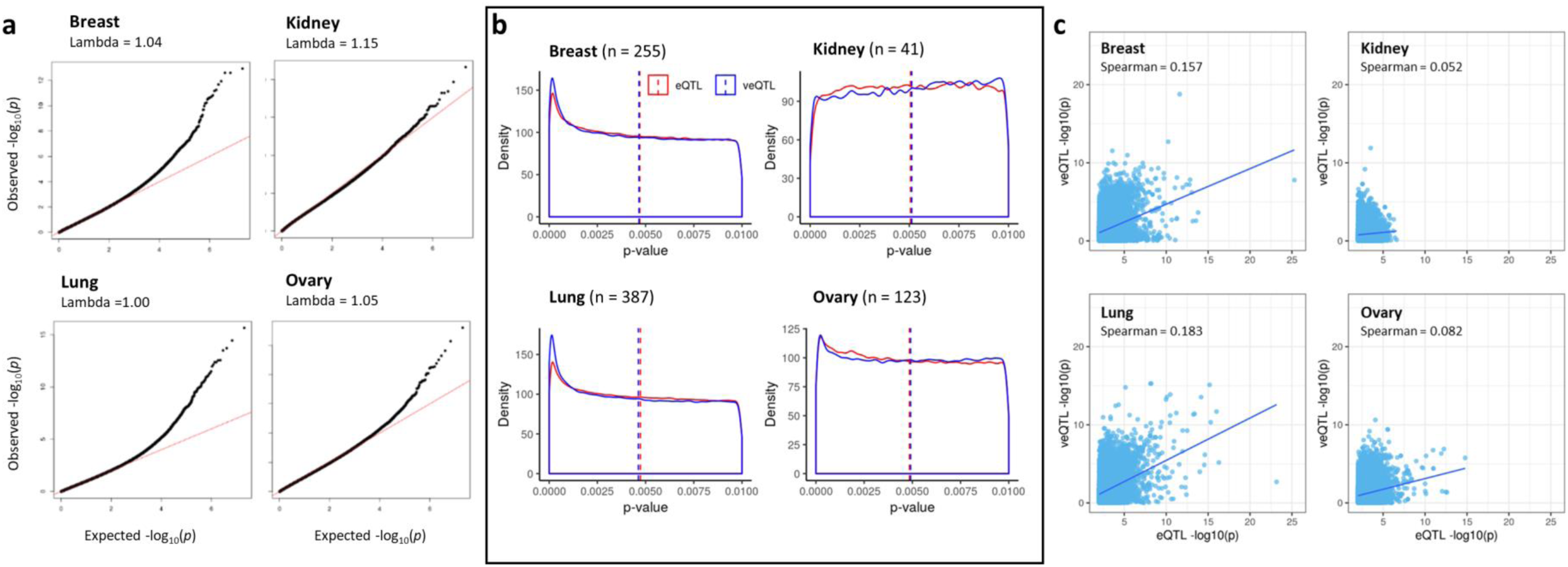
Tissue-specific performance of veQTL and eQTL analysis. a) Tissue specific q-q plots and genomic inflation factors (λ) for the associations of breast cancer risk variants and gene expression variability, with observed p-values plotted as a function of expected p-values under the null hypothesis of no association; red areas indicate the a null distribution of p values. b) Tissue specific p-value distribution for BC variants eQTLs (red) and veQTLs (blue). c) Tissue specific correlations of –log10(p) for eQTL (x-axis) and veQTL (y-axis).

Tissue specific *p*-values distributions were similar between veQTL and eQTL analyses (Figure 3b). Three tissues (breast, lung and ovary) displayed an anti-conservative distribution with a greater number of *p*-values tending towards zero. For the larger lung and breast datasets, there was a greater number of *p*-values near zero compared to ovary tissue, suggesting a greater number of tests that reject the null hypothesis of no difference in expression variability between groups. Examination of the kidney dataset demonstrated a uniform distribution of p-values, highlighting the limited effect for the selected variants for both veQTL and eQTL analysis. Variant-gene pairs were ranked according to eQTL significance and the rank correlation of *p-*values between eQTL and veQTL analysis were calculated for each tissue specific dataset. Correlations ranged from 0.052 in the kidney to 0.183 in the lung, suggesting the variant-gene ranks between veQTL and eQTL analysis are different and veQTL analysis identified a novel set of genes associated with risk variants (Figure 3c).

### Identification of potential target genes of breast cancer risk variants

The majority of breast cancer variants have no known associations with other traits, however 25 variants have previously been associated with a phenotype other than breast cancer risk (www.gwascentral.org, Supplementary Table 3). Two variants (rs11571833 and rs17879961) have been previously associated with lung cancer, while rs10069690 and rs74911261 have been associated with ovarian and kidney cancers, respectively. Interestingly, none of these variants were significantly associated with differential variability in any genes in these tissues. However, rs10069690 did have significant association with differential variability in gene expression in each of the lung and breast analysis. As the majority of the variants confer breast cancer risk only, we eliminated any veQTL that was observed in a non-breast tissue (Figure 4). Fifty-five of the 70 significant breast veQTL were observed in breast tissue only. Pathway enrichment analysis of the candidate genes associated with these breast-specific veQTL revealed hormonal biosynthetic processes and collagen fibril organisation pathways that were enriched (Figure 4**Figure 4**). The enrichment of the hormonal pathways listed in Figure 4 were driven by four genes (*CYP11B1, CYP17A1 HSD3B2* and *STAR*) all of which were associated with the risk variant rs11075995. By comparison, the 88 veQTL that were significant in lung tissue were not significantly enriched for any pathway using pathway analysis (data not shown).

**Figure 4.**
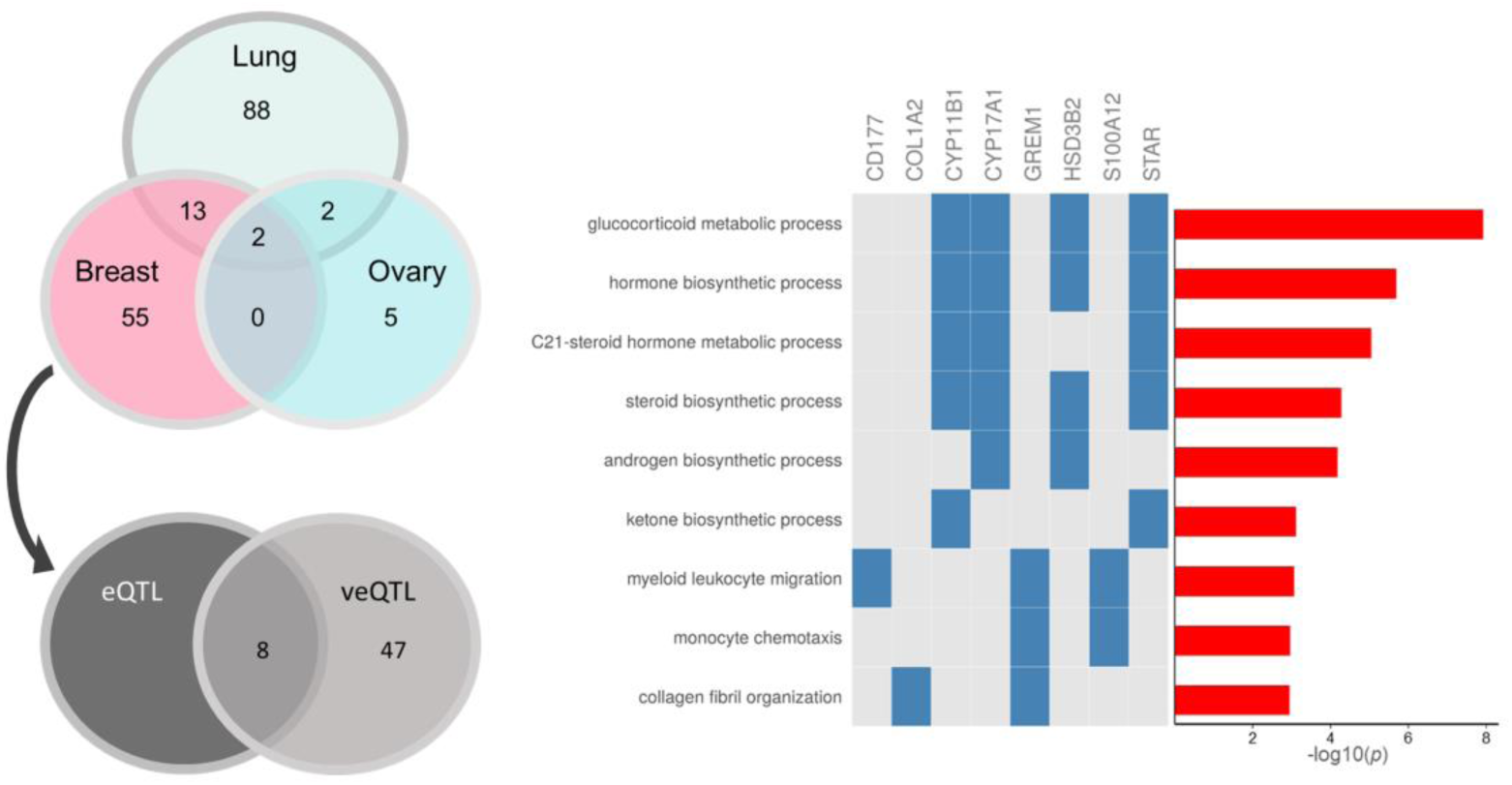
Pathway enrichment of candidate breast cancer risk genes identified through veQTL analysis. Fifty-five gene SNP pairs were observed only in breast tissues, 47 of these were veQTL but not eQTL associations. The candidate genes identified by these 47 genes were enriched for pathways involved in C21-steroid hormone metabolic process. Pathway analysis was performed in R using GO terms and using the DOSE and ClusterProfiler packages.

### rs11075995 alters expression of genes involved in C_21_ steroid synthesis

The minor allele (A) of rs11075995, which is associated with ER negative breast cancer risk, was found to be associated with increased variability in expression of four genes by veQTL analysis (Figure 5). To connect the signals of veQTL analysis with the association of breast cancer risk, we utilised the GWAS signals generated by Michailidou and colleagues on the largest meta-analysis of breast cancer risk to date and on veQTL signals generated using the GTEx data^3^. Regional plots at the rs11075995 locus for ER negative breast cancer risk associations or *trans-*veQTLs with candidate genes were visually examined to determine likely casual variants (Figure 5). Two signals were identified associated with ER negative breast cancer risk, one of which was the lead variant rs11075995 (Figure 5a). The same variants (rs11075995) produced the strongest signal for variable expression of all four candidate genes involved in the C21-steroidal pathway (Figure 5b).

**Figure 5.**
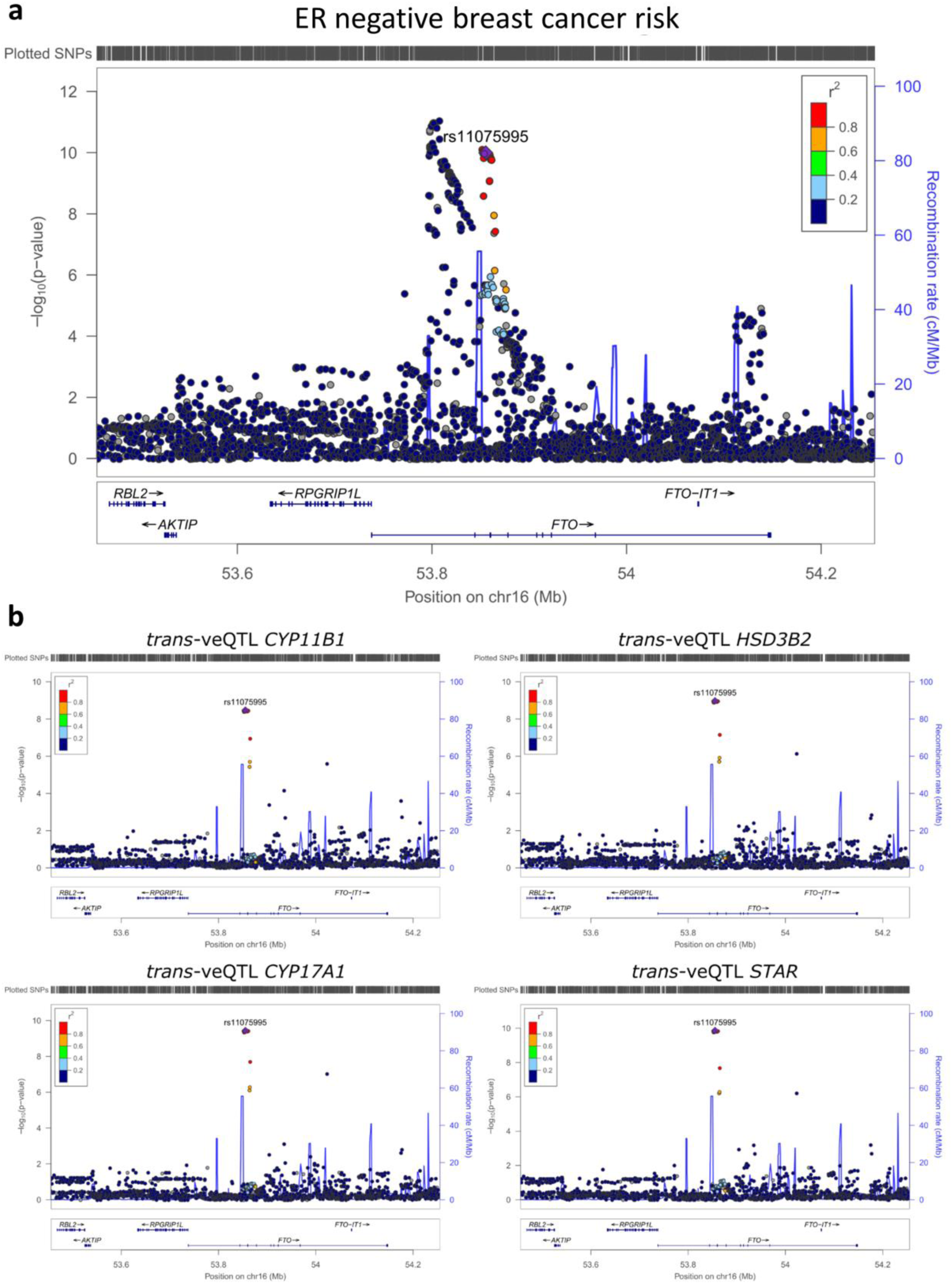
Co-localisation of ER negative breast cancer GWAS and trans-veQTL signals. a) Regional association plots for ER negative breast cancer risk for rs11075995 from Michailidou et al (2017). b) Regional association plots for trans-veQTL at rs11075995. Points indicate individual SNPs at their chromosomal location and significance (-log10(p-value)) for either GWAS (a) or trans-veQTL (b). The blue line represents the recombination rate and the colour of the points indicate the strength of the LD with rs10075995 measured as r2 in the EUR population from 1000 genomes (hg19). All plots were generated using LocusZoom.

The candidate genes (*CYP11B1, CYP17A1 HSD3B2* and *STAR*) associated with rs11075995 are all involved in the conversion of cholesterol to hormones via the C21 steroidal biosynthesis pathway (Figure 6). STAR is involved in the transportation of free cholesterol into the mitochondria where it is converted to pregnenolone. The remaining three candidate genes all code for enzymes that catalyse the conversion of multiple molecules and act in several pathways which produce different hormones (Figure 6).

**Figure 6.**
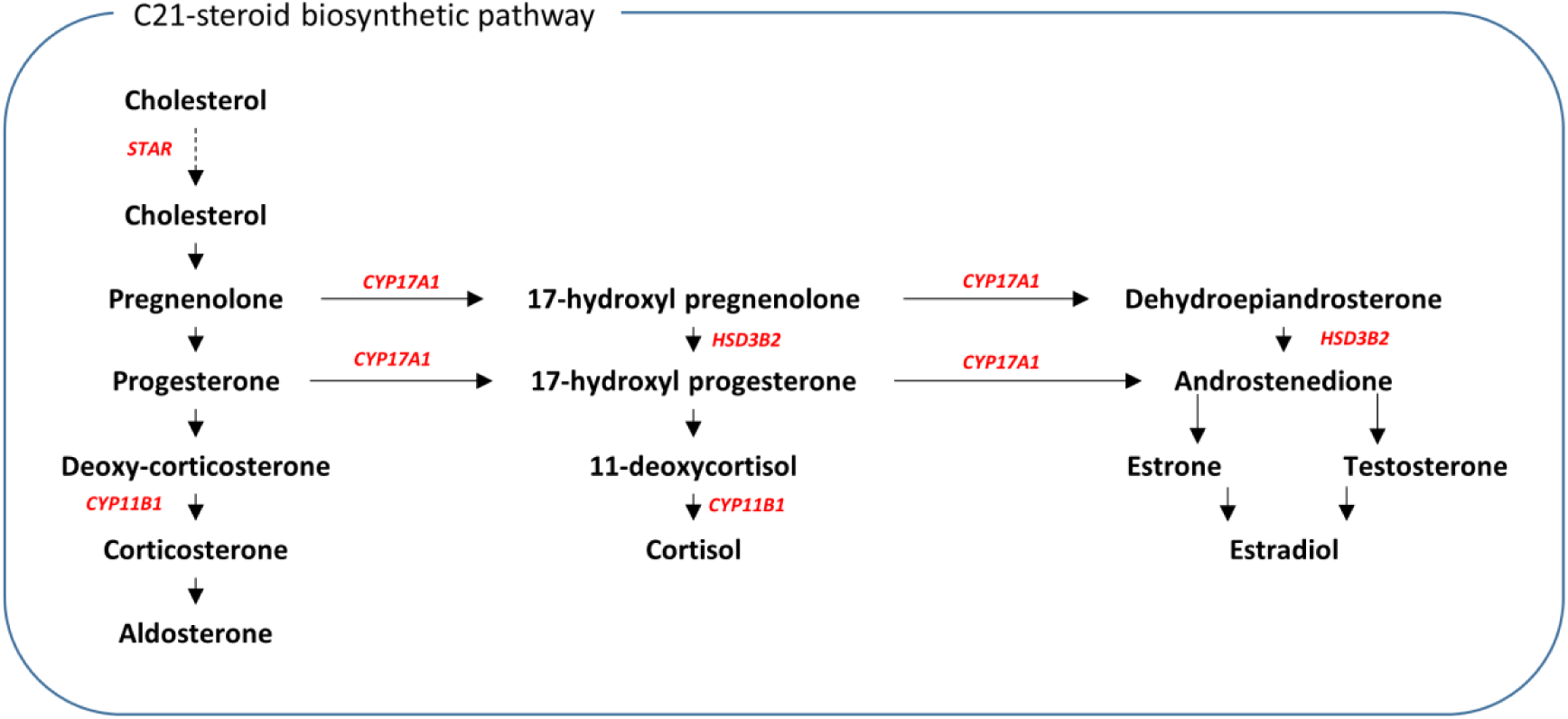
Schematic of part of C21-steroird biosynthesis pathway. Genes shown in red were associated with a significant increase in variability in individuals homozygous for the rs1105995 risk allele (A) in breast tissue (i.e. 4 of the 70 breast-derived genes from Figure 4).

The rs11075995 SNP is located in the second intron of the *FTO* gene (Figure 7)., a Fe^2+^/2-oxoglutarate-dependent oxidative RNA demethylases important in the demethylation of RNA methyladenosine (m6A)^25^. Variants in this locus are associated with increased body mass index (BMI), the mechanism of action has been linked to expression changes of the neighbouring gene *IRX3* in the human brain and in particular the hypothalamus^26,27^. Furthermore, there is conflicting evidence of rs11075995 association with breast cancer risk. Recent studies identified a loss of breast cancer risk association after adjusting for BMI^28^. However, Garcia-Closas and colleagues tested the association with ER negative breast cancer risk after adjusting for BMI and observed no change^29^. We therefore explored the effects of the rs11075995 on the expression of both *FTO* and *IRX3* in breast tissue. Neither *FTO* nor *IRX3* had significant breast eQTL or veQTL associations with rs11075995. However, *FTO* (*p* = 0.05), and not *IRX3* (*p* =0.29), had decreased expression in the homozygous minor allele individuals in breast tissue (Figure 7b, Supplementary Figure 1).

**Figure 7.**
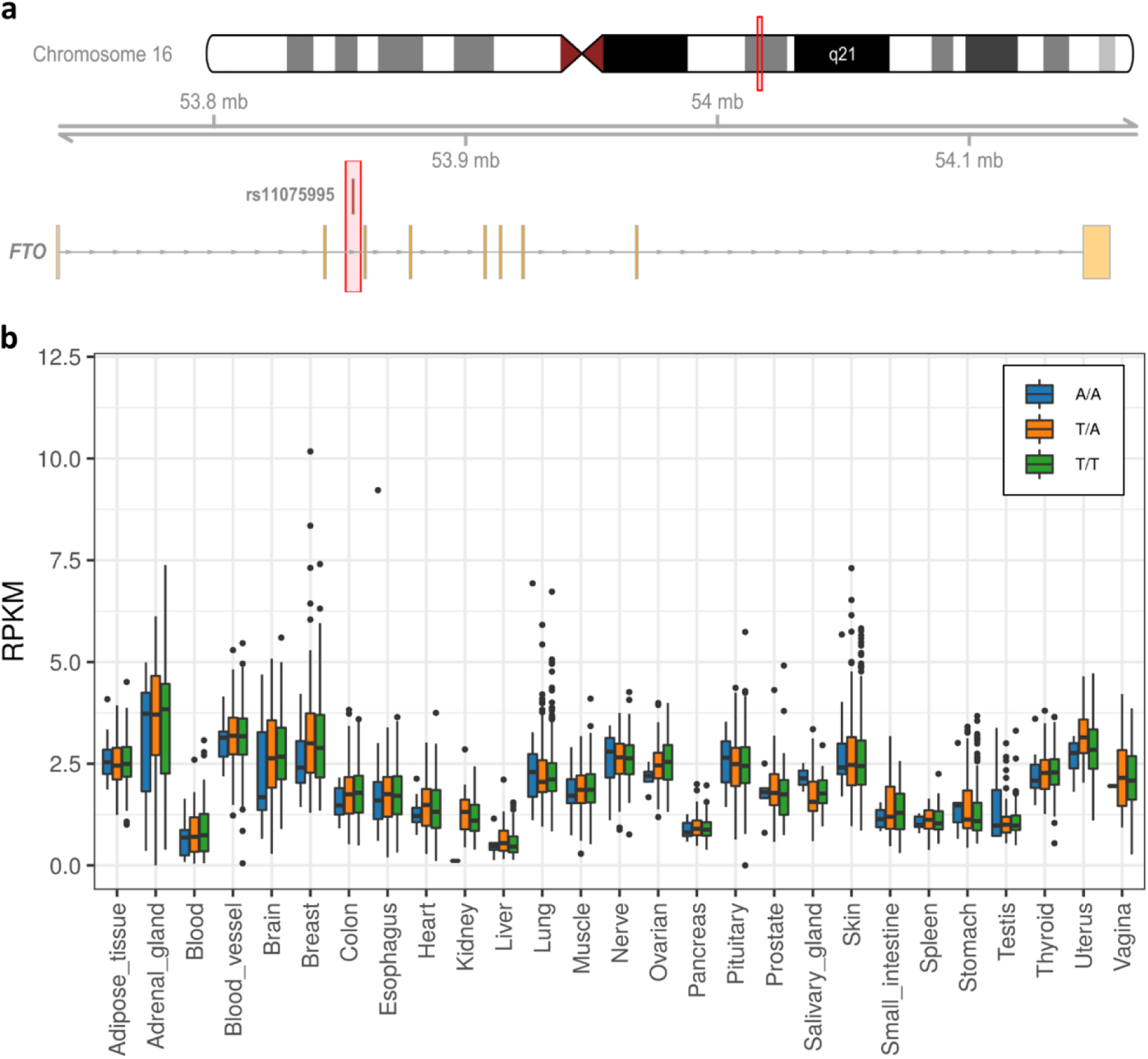
cis-effects of rs11075995 minor allele and FTO expression. a) Ideogram and chromosomal location of the rs11075995 variant within in the intron FTO gene. b) Tissue specific expression of FTO stratified by genotypes at the rs11075995 location. T/T homozygous major allele (Green), A/T heterozygous (Orange), A/A homozygous minor allele (Blue).

## DISCUSSION

Tissue-specific veQTL datasets were generated for breast cancer variants in four normal tissues dataset acquired from GTEx. To predict candidate genes involved in breast cancer risk, significant (*p* < 5×10^−8^) veQTL unique to breast tissue were considered. This approach identified 60 candidate genes that were associated with 27 variants. The majority of significant veQTL were class I and displayed a homozygous recessive like phenotype (Figure 2). Furthermore, veQTL analysis identified distinctly different genes compared to eQTL analysis (Figure 3). Although, 30% of class II breast veQTL were also eQTL, highlighting a small subset of genes that had both changes in mean expression and variability associated with minor allele dosage.

Pathway analysis of the 60 candidate genes found several hormonal biosynthetic pathways enriched along with monocyte chemotaxis and collagen fibril organisation (Figure 4). The enrichment of the hormonal biosynthetic pathway was driven by the presence of four genes (*CYP11B1, CYP17A1, HSD3B2* and *STAR*) all of which were variable in association with the risk allele of rs11075995. Furthermore, rs11075995 produced the strongest signal for variable expression for all four candidate genes and was the most likely casual variant (Figure 5).

Breast cancer development has been associated with exposure to steroid hormones^30^. These hormones are typically synthesised in non-breast tissues (e.g. ovary and adrenal gland) and are secreted into the circulating system to act on distant tissues (e.g. breast). The activation of local hormone biosynthesis, associated with the risk allele of rs11075995, through the metabolism of cholesterol to pregnenlone may lead to greater exposure and/or hormone imbalance in breast tissues, which may drive tumourigenesis. Local steroidogenesis and ultimately production of androgens has been observed in androgen independent advance prostate cancers^31^. In prostate cancer, the local production of androgens may explain the development of hormonal treatment resistance in late-stage prostate cancers.

Summary statistics of GWAS signals obtained through GWAS central (www.gwascentral.org) identified significant associations of rs11075995 with overall and ER negative breast cancer risk and with body mass index (Supplementary Table 3). No other trait was reported to be associated at *p* < 0.001 with rs11075995. BMI is a known dose-dependent risk factor for developing breast cancer in post-menopausal women^32^. Interestingly, breast cancer risk association studies that have adjusted for BMI have demonstrated a dependence for variants at the rs11075995 locus on BMI status^28^. However, an independent relationship was described for ER negative breast cancer risk and BMI for rs11075995^29^, suggesting that variants in the same locus may have disease-specific risk profiles.

The variant rs11075995 is located in intron 2 of the *FTO* gene. Interestingly, we observed a marginally significant decrease in *FTO* (p = 0.05) expression in breast tissue associated with individuals homozygous for the rs11075995 risk allele. FTO is involved in demethylation of RNA adenosine (m6A). Methylated adenosine are post-transcriptional modifications which signals RNAs for processing, including degradation and splicing^34^. The four genes associated with rs11075995 all have the m6A target site (GGACU). RNA variability may occur due to dysregulation of these pathways (mRNA degradation and splicing) in response to decreased *FTO* expression

Variants in intron 1 and 2 of *FTO* have been strongly associated with obesity and changes in BMI^26,33^, however these variants act on the expression of the neighbouring gene *IRX3* in the hypothalamus region of the brain^27^. Iroquois homeobox protein 3 (IRX3) is a highly conserved transcription factor typically expressed during neural development^35^. The role of *IRX3* in obesity is yet to be fully elucidated with conflicting reports of body mass associated to deficient *Irx3.* Smemo et al., described a 30% increase in body weight of *Irx3-*deficient mice^27^. While in contrast the partial depletion of *Irx3* through a lentiviral system resulted in mice with greater body mass^36^.

Intriguingly, both *IRX3* and *FTO* are highly expressed in the hypothalamus, a region of the brain important to hormonal regulation^27,36^. It is unknown whether risk variants, for either BMI or breast cancer, directly disrupt the regulation of hormonal control in the hypothalamus. Furthermore, it is unclear what effect *IRX3* expression would have on breast cancer risk and whether any effect would be independent of the risk attributed to obesity alone. A better understanding of the downstream transcriptional targets of IRX3 may identify pro-tumourgeneic pathways.

Our results are consistent with the hypothesis that different variants in the *FTO* locus may be associated with tissue-specific hormonal control and subsequently different pathologies. Consequently, we would expect differences in the regulation of C21 hormones in breast tissue for the different rs11075995genotypes. Furthermore, candidate genes identified through veQTL analysis require functional validation. A major challenge with assessment of intra-sample gene expression variability is the limitation of single-point ‘grind and bind’ approaches. However, approaches such as RNA hybridisation *in situ* and single cell RNA-sequencing do provide the ability to detect expression variability. It is of further importance to derive the mechanism of variability which may be driven by interaction of genotypes with exposures or epistasis.

In summary, breast cancer risk variants are associated with variable expression of candidate breast cancer susceptibility genes. These included genes involved in hormonal biosynthetic pathways that are associated with a single variant (rs11075995). To our knowledge, this is the first time gene expression variability has been used to identify candidate cancer susceptibility genes.

---

**Supplementary Figure 1.**
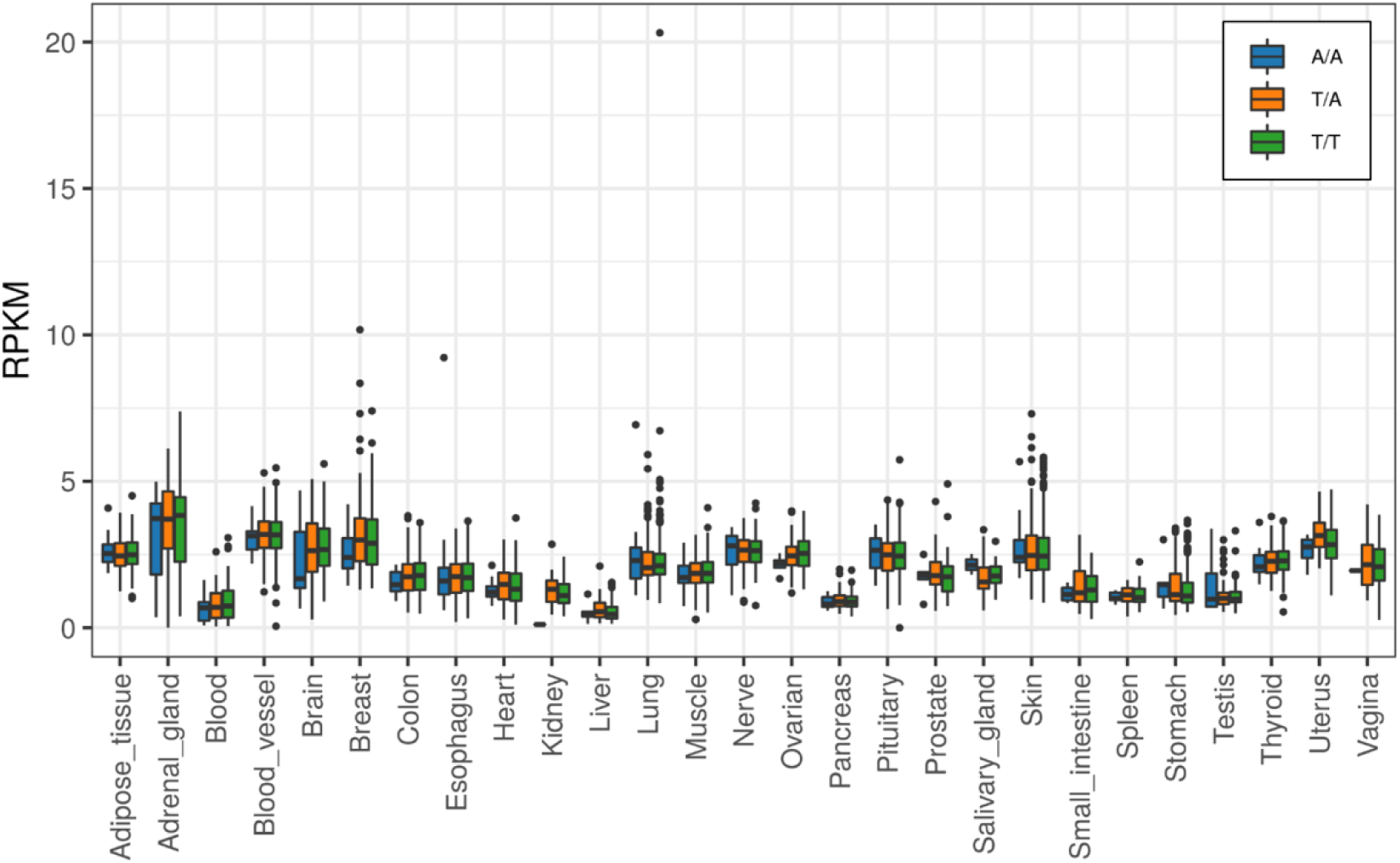
Tissue specific expression of IRX3 stratified by genotypes at the rs11075995 location. T/T homozygous major allele (Green), A/T heterozygous (Orange), A/A homozygous minor allele (Blue).

